# Imaging biological samples by integrated differential phase contrast (iDPC) STEM technique

**DOI:** 10.1101/2021.07.20.453030

**Authors:** Xujing Li, Ivan Lazić, Xiaojun Huang, Li Wang, Yuchen Deng, Tongxin Niu, Dongchang Wu, Maarten Wirix, Lingbo Yu, Fei Sun

**Affiliations:** Center for Biological Imaging, Institute of Biophysics, Chinese Academy of Sciences, Beijing 100101, China; National Key Laboratory of Biomacromolecules, CAS Center for Excellence in Biomacromolecules, Institute of Biophysics, Chinese Academy of Sciences, Beijing 100101, China; Thermo Fisher Scientific, Materials and Structural Analysis, Eindhoven, The Netherlands; University of Chinese Academy of Sciences, Beijing, China; Physical Science Laboratory, Huairou National Comprehensive Science Center, No. 5 Yanqi East Second Street, Beijing 101400, China; Bioland Laboratory (Guangzhou Regenerative Medicine and Health Guangdong Laboratory), Guangzhou, Guangdong 510005, China

**Keywords:** Biological sample, Contrast, Integrated differential phase contrast, Scanning transmission electron microscopy, Thick section

## Abstract

Scanning transmission electron microscopy (STEM) is a powerful imaging technique and has been widely used in current material science research. The attempts of applying STEM into biological research have been going on for decades while applications have still been limited because of the existing bottlenecks in dose efficiency and non-linearity in contrast. Recently, integrated differential phase contrast (iDPC) STEM technique emerged and achieved a linear phase contrast imaging condition, while resolving signals of light elements next to heavy ones even at low electron dose. This enables successful investigation of beam sensitive materials. Here, we investigate iDPC-STEM advantages in biology, in particular, chemically fixed and resin embedded biological tissues. By comparing results to the conventional TEM, we have found that iDPC-STEM not only shows better contrast but also resolves more structural details at molecular level, including conditions of extremely low dose and minimal heavy-atom staining. For thick sample sections, iDPC-STEM is particularly advantageous. Unlike TEM, it avoids contrast inversion canceling effects, and by adjusting the depth of focus, fully preserves the contrast of relevant features along with the sample. In addition, using depth-sectioning, iDPC-STEM enables resolving in-depth structural variation. Our work suggests that promising, wide and attractive applications of iDPC-STEM in biological research are opening.

## INTRODUCTION

The invention of electron microscope has been a great millstone for human to observe the micro-world and investigate the nature at nano and atomic scale. Various electron microscopy (EM) techniques, e.g., transmission electron microscopy (TEM), scanning electron microscopy (SEM), scanning transmission electron microscopy (STEM) and so on, have been developed in the past seventy years. Vast amount of structural and dynamics information from various diverse materials have been obtained by electron microscopy. However, the application of EM to study biological specimen has been delayed for many years due to four major limitations. First, the vacuum of specimen chamber of electron microscope is in conflict with the need of water for live biological sample; second and more important, biological samples are highly sensitive to electron radiation (Baker and Rubinstein, 2010; Glaeser, 1971); third, the thickness of interest of biological samples is normally beyond the suitable penetration depth of the electron; and fourth, biological samples are mainly consisting of light elements (C, H, O, N, P and S), resulting a relative low contrast with normal EM techniques.

To overcome these challenges for biological TEM, large number of efforts have been performed in the past. The first attempt was to develop chemical fixation and dehydration protocols and embed biological sample into resin by trying to keep close to native state. The ultrathin section of resin embedded biological sample can be imaged directly by normal TEM. Heavy metal staining (osmium oxide, uranyl acetate or lead citrate) was always used to enhance the TEM contrast of the resin section. With this procedure, biologists obtained great achievements by discovering fruitful information of cellular ultrastructures in the last century. The second attempt was to develop cryo-fixation protocol and the technique of cryo-electron microscopy (cryo-EM) (Adrian et al., 1984; Dubochet et al., 1988). The cryo-vitrified biological samples keep their most natural state and can be imaged to atomic resolution by cryo-EM (Nakane et al., 2020; Yip et al., 2020). Nowadays, cryo-EM has become one of the major biophysical techniques to study 3D structures of bio-macromolecules, giving a great impact on the life science research.

Introducing other EM techniques that are already available for materials science research into biological research has been always welcome with the aim to compensate for the current biological TEM techniques shortcomings and provide new useful information of biological samples. As a fully proved successful technique in characterizing materials at micro, nano, and atomic scales for a vast variety of different samples, STEM technique has been widely utilized and nowadays become one of the major high-resolution imaging tools in materials science. STEM imaging mode is obtained with an electron beam that is finely focused to a small probe at the plane of the specimen and then scanned over it. Depending on the convergence angle of the beam, the depth of focus of STEM can easily be altered to become longer than a given sample thickness. The probe scans the sample point by point, and transmitted electrons are collected and counted for each point by STEM detectors in the far field. The electrons scattered to different angles can produce different images. The brightfield (BF) detector collects electrons scattered within the convergent angle of the beam, whereas annular darkfield (ADF) detectors collect electrons scattered outside of the beam convergent angle regions, often to relatively high angles (HA) of ADF (see various literatures summarized in text books (Reimer and Kohl, 2008; Spence, 2013) and recent analytical in depth analysis (Bosch and Lazić, 2015; Lazić and Bosch, 2017)).

Applying STEM in biological research was originated only couple of decades ago (Colliex and Mory, 1994). Compared to TEM, STEM generate variety of signals, which can be used for biomacromolecule mass quantitative measurement, elemental mapping, and multi-signal imaging (Aronova and Leapman, 2012; Colliex and Mory, 1994; Sousa and Leapman, 2012). With the Z^2^ -contrast feature of (HA)ADF-STEM favoring heavy elements (Bosch and Lazić, 2015, Lazić and Bosch, 2017), STEM has the unique advantage of visualizing nanogold particles (1.4 nm in diameter) embedded in the biological resin section (Sousa et al., 2007), which is very important for the immune-EM research. The beneficial aspect of STEM in electron tomography of resin embedded biological section has also been investigated (Aoyama et al., 2008; Hohmann-Marriott et al., 2009; Sousa and Leapman, 2012; Yakushevska et al., 2007). Compared to TEM tomography, STEM tomography could generate a tomogram with better contrast and better signal-to-noise ratio (SNR) (Yakushevska et al., 2007). In addition, more importantly, some STEM techniques have a large depth of focus, no oscillation of contrast transfer function (CTF) (e.g. HAADF-STEM) and can achieve dynamic focusing, which endows STEM tomography a unique advantage of studying thick (1 μm) biological sections (Aoyama et al., 2008). Considering the multi-scattering effect of electrons, the resolution and contrast of STEM tomography of thick section could be further improved by changing dark field to bright field (Hohmann-Marriott et al., 2009). The attempts of applying STEM tomography to frozen-hydrated biological sample were also performed in recently years (Rez et al., 2016; Trepout, 2020; Wolf and Elbaum, 2019; Wolf et al., 2014), which turned to be useful to detect metabolic ions in biomacromolecules (Elad et al., 2017), study mitochondrial calcium stores physiology (Wolf et al., 2017) and dynamics of ferritin crystallization (Houben et al., 2020). Besides collecting tilt series like conventional electron tomography, STEM offers another possibility of tomography by depth section (Bosch and Lazić, 2019; Trepout et al., 2015).

Although many above efforts of trying STEM imaging in biological applications have been performed, this technique has not yet been widely used in the current biological research. Although (HA)ADF-STEM could yield a good, interpretable contrast, the number of electrons collected in the dark field is several orders of magnitude smaller than that in the bright field, effectively making (HA)ADF-STEM extremely dose inefficient with bad signal to noise ratio (SNR) and not suitable for dose sensitive biological samples. Furthermore, the contrast of (HA)ADF-STEM imaging is proportional to the square of elemental Z value (Lazić and Bosch, 2017), causing the biological light elements (C, H, O, N, etc.) to be less visible when imaged next to heavier ones (Si, P, S etc.) or invisible next to heavy ones (Ga, Au, Os, Ur, Pb, etc.). The image contrast of (annular) bright field ((A)BF) STEM is nonlinear, difficult to interpret, and the CTF of (A)BF-STEM also suffers from the shortcoming of oscillations (Lazić and Bosch, 2017) the same way TEM does, which is why it does not show any superior advantage over the conventional TEM.

An appropriate STEM imaging technique for biological samples should generate linear contrast with positive definite CTF, including all frequencies and using all possible available electrons. Such a technique has been introduced, called Integrated Differential Phase Contrast STEM (iDPC-STEM) (Lazić et al., 2016; Yucelen et al., 2018). The iDPC-STEM is based on the detection of center of mass (COM) or center of illumination of the beam in the detector plane, which can be effectively accomplished already with 4-quadrant segmented detectors. The COM signal linearly maps the electric *vector* field of the sample and an integration (inverse gradient) of this vector map yields a *scalar* electrostatic potential field of the sample, resulting in a final image with the linear phase contrast. Theoretical calculations proved the CTF of iDPC-STEM is definite positive without any oscillations, and frequency transfer includes low frequencies. Therefore, using iDPC-STEM technique, light and heavy elements can be successfully imaged together (Gauquelin et al., 2017; Nahor et al., 2018; Song et al., 2019; Yucelen et al., 2018), including hydrogen atoms (de Graaf et al., 2020). In addition, the integration step of iDPC-STEM makes a natural suppression of the non-conservative part of the noise, i.e. only conservative field (here electric field) can be integrated. Due to this, a superior SNR is obtained (Lazić and Bosch, 2017; Lazić et al., 2016) and a low dose imaging is enabled for the carbon-based and other beam sensitive materials like zeolites (Shen et al., 2020b; Shen et al., 2021) and metal organic frameworks (MOFs) (Shen et al., 2020a) with quality better than the one offered by conventional TEM.

Although iDPC-STEM has been successfully and widely used in material science as listed above, its advantages for biological samples have not been explored yet. In this work, for the first time, we demonstrate ability of iDPC-STEM to image chemically fixed and resin embedded biological tissue and cell samples. We found that, compared to TEM images, iDPC-STEM images could not only show better contrast but also resolve more cytosolic details in molecular resolution on the conditions of either low dose or minimal heavy atom stains. Furthermore, for the thicker sections (400-600 nm thickness), iDPC-STEM is superior and can resolve more structural details where the conventional TEM suffers from contrast inversion canceling effects (the defocus gradient induced signal fading due to thickness) and fails. For completeness, an initial trial of depth-sectioning of a thick biological section has been performed showing capability to resolve in-depth structural information.

## MATERIALS AND METHODS

### Theoretical consideration of the beam convergence semi-angle

A focused electron beam in STEM, called the probe, takes different shapes along the direction of propagation depending on electron energy, beam convergence semi-angle and aberrations introduced by forming condenser lenses. For most practical purposes, all aberrations except spherical aberration can be corrected to acceptable values by well-designed cylindrically symmetric column, proper alignment and stigmation correction. The regular STEM probe along the electron propagation shows a spindle-like shape (Figure 1a). Most intensity of the probe is concentrated within the region of the spindle, which interacts the strongest with the sample electric field and contributes dominantly to the signal at the detector and, therefore, the contrast of STEM images (Bosch and Lazić, 2019). With fixed spherical aberration, the size of STEM probe depends directly on the convergence semi-angle α that is determined by the condenser lens aperture. The width of the probe determines the x-y resolution of STEM imaging, and the length of the probe determines the depth of focus of STEM imaging, the z resolution in depth-sectioning mode. Within the critical condition (see description below) the larger the convergence semi-angle, the smaller the probe width and the probe length are, and vice versa. For a better resolution and reduced depth of focus, a larger convergence semi-angle is needed (Table 1).

**Figure 1.**
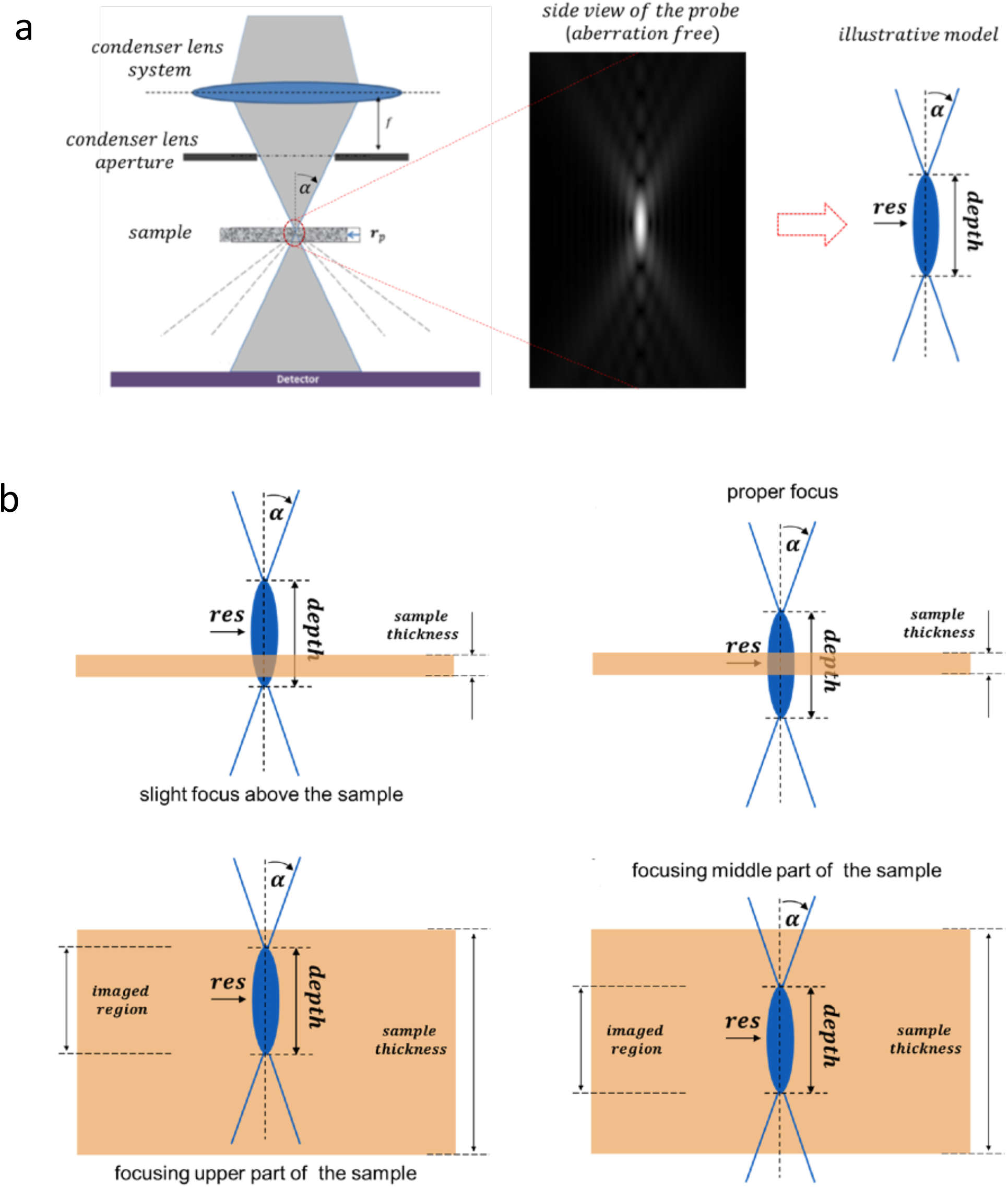
Theoretical diagram of STEM imaging. (**a**) Cross-section of STEM imaging system and the STEM probe. Enlarged vertical cross-section of the probe (middle) and illustrative probe model (right) are shown with the key parameters (convergence semi-angle α, resolution and depth) of the probe indicated. (**b**) Two main situations of STEM imaging in relation to the depth of focus vs. the sample thickness. Top: the depth of focus is larger than the sample thickness, the best in-focus image is obtained by placing the position of the sample just at the middle of the spindle-shaped probe (disc of least confusion). Bottom: the depth of focus is shorter than the sample thickness, different parts of sample are imaged with different focal position of the probe and the depth-sectioning can be performed.

**Table 1.** The dependency of STEM resolution and depth of focus with the convergence semi-angle for the 200 keV electron (wavelength ***λ*** = **2**. **508 *pm***)*.

| Convergence semi-angle $\alpha$ [mrad] | 3 pixel resolution at proper pixel size [Å] | Ultimate resolution $\lambda/(2\alpha)$ [Å] | Proper pixel size for ultimate resolution (Nyquist criterion) [Å] | Depth of focus $2\lambda/\alpha^2$ [nm] |
| --- | --- | --- | --- | --- |
| 1 | 18.7 | 12.5 | 6.2 | 5016 |
| 2 | 9.4 | 6.3 | 3.1 | 1254 |
| 3 | 6.3 | 4.2 | 2.1 | 555 |
| 4 | 4.6 | 3.1 | 1.5 | 312 |
| 5 | 3.7 | 2.5 | 1.2 | 200 |
| 6 | 3.1 | 2.1 | 1.0 | 139 |
| 7 | 2.7 | 1.8 | 0.9 | 102 |

There are two STEM imaging modes used in this work (Figure 1b). The first mode involves the depth of focus longer than the sample thickness. In this mode (Figure 1b, top) the sample can be considered thin with a high level of accuracy (Bosch and Lazić, 2019), and the image formation can be fully described with the thin sample STEM theory (Bosch and Lazić, 2015). In this mode the in-focus image is obtained when the sample is located at the position of the beam central cross-over (the disc of least confusion), while a slightly defocused image is formed when the central position of the probe is apart above (over focus) or below (under focus) the sample. For iDPC-STEM and ADF-STEM the in-focus image will have the best contrast. Unlike that, for (A)BF-STEM the most contrast will occur out of focus, for defocus value called Scherzer defocus (Kirkland, 2010). The second imaging mode involves the depth of focus shorter than the sample thickness (Figure 1b, bottom). In this mode the sample cannot be considered thin, and the depth-sectioning of the sample takes place (Bosch and Lazić, 2019). By changing defocus, one could cover and image the full 3D volume of the sample with the resolution in-depth determined by the length of the beam depth of focus region (de Jonge et al., 2010).

When the spherical aberration (Cs) of the condenser system is not corrected the smallest probe is defined with a critical convergence semi-angle, determined with *α*_*max*_ = 1.34 *λ*/(*C*_*S*_*λ*^3^)^1/4^, where *λ* is relativistic wavelength of the electron, see analysis by Kirkland (Kirkland, 2010) (Sec. 3.5.1). For the convergence semi-angles smaller than this critical value the probe is practically the same in shape and size as for corrected systems, while further increasing the convergence angle results in a dramatically deteriorate shape of the probe in all three directions (Figure S1). In this work, we used an aberration non-corrected microscope and carefully set up the convergence semi-angle smaller than this critical value.

### Electron microscope set-up and imaging parameters

In this work, all the electron microscopy images (both TEM and STEM) were acquired using FEI Talos F200C (Thermo Fisher Scientific), an electron microscope with a two-condenser lens, spherical aberration non-corrected (Cs = 2.7 mm) system, operated at 200 kV. For the conventional TEM imaging, an equipped 4*k x* 4*k* Ceta CMOS camera and an objective aperture (100 μm) were used. For STEM imaging, standard BF and HAADF detectors were used for usual (A)BF and ADF modes, as well as standard segmented 4-quadrant detector for iDPC-STEM mode, and condenser apertures (20 μm, 50 μm and 70 μm). All iDPC-STEM and TEM images were taken and processed with the Velox software. For comparison of TEM and STEM images, the pixel size for STEM has been selected as close as possible to the one applied in TEM.

The critical value of the convergence semi-angle of our electron microscope system before probe starts deteriorating, based on formula given above and *C*_*S*_ = 2.7 *mm*, is *α*_*max*_ = 7.4 *mrad*. Therefore, the dependency of the STEM resolution and depth of focus with respect to convergence semi-angles smaller than this critical value have been computed and shown in Table 1. With available apertures and condenser lens modes the two convergence semi-angles of 2.58 mrad (the corresponding depth of focus is 751 nm) and 6.43 mrad (the corresponding depth of focus is 121 nm) have been selected for STEM imaging in this work.

### Electron dose calculation

In TEM imaging, the total dose in [*e*/Å^2^] units is calculated as total number of electrons (in [*e*] units) passed per square area (in [Å^2^] units) during exposure time,

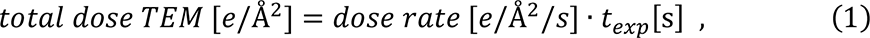

where the dose rate is beam current per unit area (beam current density). The beam current, usually calibrated and read in [*pA*] = [*pQ*/*s*] is converted into [e/s] units, using 1.00 [*Q*] = 1.00/(1.60 · 10^−19^)[*e*] = 6.24 · 10^18^[*e*], before dose rate is computed, yielding 1.00 [*pA*] = 6.24 · 10^6^ [*e*/*s*]. The TEM exposure time *t*_*exp*_ is measured in seconds.

In STEM, the total dose in [e/Å^2^] units is calculated as total number of electrons passed the total field of view (FOV) area (full image area) after the full scanning frame is finished. This is the same for all type of STEM including iDPC-STEM, yielding

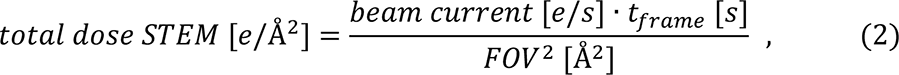

where the beam current, usually calibrated and read in [*pA*] = [*pQ*/*s*] is converted into [*e*/*s*] units, as above. The total scanning frame time *t*_*frame*_ is measured in seconds, while the total image field of view is given in [Å]. The full frame time *t*_*frame*_ is the time spent per pixel, called the dwell time *t*_*dwell*_, multiplied by the number of pixels visited by the beam. Assuming that *N x N* image is acquired, *t*_*frame*_ = *N*^2^ · *t*_*dwell*_ and *FOV* = *N* · *pixel size*, which further yields more practical formula

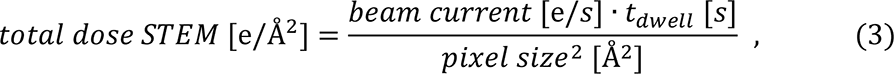

Where number of pixels simply cancels out.

The *dwell times* are normally many orders of magnitude smaller than the *frame times* (e. g. for 1*k x* 1*k* image that is one million times smaller) and, therefore, naturally measured in [*μs*]. Beam current is, for practical cases in biology, measured in [*pA*]. Taking this into account the above formula, Eq. (3), further becomes

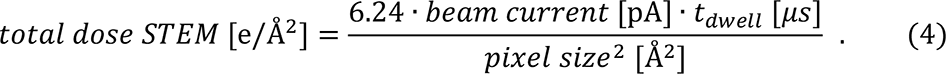

With a quick computation check we see that 1 *pA* beam current, 1*μs* dwell time with a scanning pixel size of 2.5 Å, produces the total dose of 1.0 *e*/Å^2^.

### Preparation of resin embedded tissue sample

Two samples, one for the human hepatocellular carcinoma cell (HepG2) and another for the mouse testis tissue, were selected in this work to prepare resin embedded sections. Two sample preparation protocols, one with regular heavy atom staining (Method 1) and second with minimal staining (Method 2) were performed, respectively (Figure S2).

For the protocol of regular heavy atom staining (Method 1), after euthanasia of a mouse at 4°C, tissues were quickly dissected and fixed in the fixative solution containing 2.5% glutaraldehyde for 48 hr. For HepG2 cells, the cells were collected and washed 3 times with PBS and fixed with 2.5% glutaraldehyde at 4℃. All samples were washed with PBS and further fixed with the buffer containing 1% OsO4 for 2 hr. The chemically fixed sample were dehydrated using graded ethanol solutions from 20% to 100% and embedded in Epon 812 resin (SPI Supplies of Structure Probe, Inc.). Ultrathin sections of resin embedded tissue sample were obtained using ultramicrotome UC7 (Leica Microsystems) and further stained using 2% uranyl acetate for 30 minutes and then lead citrate for 10 minutes.

For the protocol of minimal heavy atom staining (Method 2), the procedures from tissue dissection/cells collection, fixation and dehydration to resin embedding and ultrathin sectioning were exactly same as Method 1. The only difference is that there was no further post staining on the ultrathin sections.

### Experimental set-up to investigate iDPC-STEM technique

In this work we set up four kinds of experiments to investigate iDPC-STEM imaging technique and make comparisons with the conventional TEM. First, we investigate image contrast of the regularly stained tissue sections by changing electron dose. We determine the required doses for iDPC-STEM and TEM, respectively, for which the optimal contrast was obtained. Then, while keeping the electron dose at which the conventional TEM imaging shows a good contrast (for the regularly stained sample), we switch to the minimally stained samples and compare the contrast between TEM and iDPC-STEM again. Next, we increase the thickness of the section from commonly used 70 nm to 200 nm, 400 nm, even up to 600nm and compare the contrast and image quality between TEM and iDPC-STEM again. Finally, in-depth structural information acquisition of the thicker section with iDPC-STEM depth-sectioning is demonstrated.

### Quantitative measurement of contrast

To qualitatively compare the contrast in the images obtained using different techniques and conditions, a method of line profile over the area of interest is used (Figure S3). First, the images are normalized according to the pixel variance (step 1). Then the line profile is generated at the region of interest using Gatan Digital Microscopy software (step 2). Finally, the contrast is computed as difference between maximum and minimum intensity along the profile, normalized to the maximum (step 3).

## RESULTS

### iDPC-STEM contrast vs. TEM contrast with respect to electron dose

We make the first comparison between iDPC-STEM and TEM using the regularly prepared thin (70 nm) section (Method 1) of resin embedded HepG2 cell. We set the convergence semi-angle of iDPC-STEM to a higher value, 6.43 mrad in this experiment. The corresponding depth of focus is 121 nm (Table 1), which is larger than the thickness of the section, while resolution down to 3Å is, in principle, possible. Thus, we followed the first imaging mode of STEM (Figure 1b) and optimized the z-position of the section to get the sharpest iDPC-STEM images with the best contrast. The original pixel size of iDPC-STEM images was set to 1.05 nm and then resized to match the pixel size of TEM images by FFT cropping algorithm. For TEM imaging, we set up an appropriate magnification to image the same region of the section with a similar field of view (FOW) and the final pixel size on the camera was 1.50 nm. We used a big defocus value of ∼7 μm in TEM imaging to compensate the attenuation of CTF at the low frequency and achieve a good contrast.

We kept the electron dose for each compared pair of iDPC-STEM and TEM images exactly the same and took a series of micrographs with different electron doses starting from 1 e/Å^2^, 3 e/Å^2^, 6 e/Å^2^, 10 e/Å^2^, 20 e/Å^2^ up to 40 e/Å^2^ (Figure 2a). At the relatively low dose conditions (1 e/Å^2^, 3 e/Å^2^, 6 e/Å^2^ and 10 e/Å^2^), by qualitative examination iDPC-STEM imaging produces better micrographs with better contrast and better SNR. The structural details of organelle membranes and, especially, the fine structures within the mitochondrial inter-membrane space, are clearly visualized and resolved within iDPC-STEM micrographs, while not in the corresponding TEM images. We stress here the extreme case, using electron dose of only 1 e/Å^2^, where the conventional TEM does not generate enough contrast to resolve the ultrastructure of mitochondria, while iDPC-STEM resolves the mitochondrial double membrane and crista.

**Figure 2.**
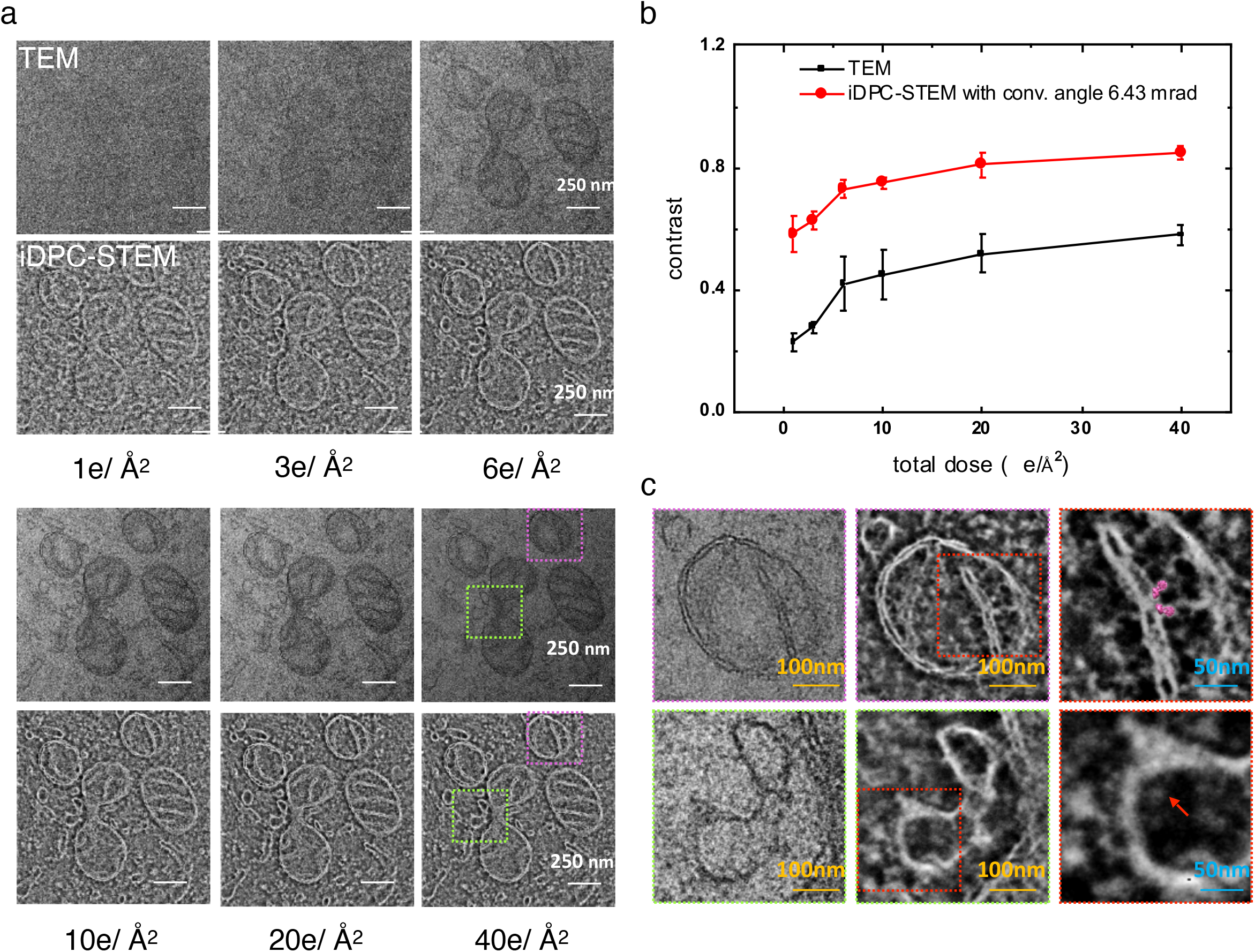
Comparison between TEM and iDPC-STEM imaging at different electron dose. (**a**) TEM and iDPC-STEM micrographs acquired on 70 nm section of HepG2 cell with the total electron doses of 1 e/Å^2^, 3 e/Å^2^, 6 e/Å^2^, 10 e/Å^2^, 20 e/Å^2^ and 40 e/Å^2^, respectively. The pixel size for TEM is 1.50 nm. The original pixel size of iDPC-STEM was 1.05 nm, resized to 1.50 nm by FFT cropping algorithm. The convergence semi-angle of iDPC-STEM is 6.43 mrad. (**b**) Quantitative comparison of contrast between TEM and iDPC-STEM at different exposure doses. The contrast of each micrograph was measured using the line profile method (see Figure S3). For each data point, the measurement was repeated three times using different profile lines at different regions. (**c**) Zoom-in comparison between iDPC-STEM and TEM images at the dose of 40 e/Å^2^. Regions of both iDPC-STEM and TEM images marked by the purple/green rectangles in (a), representing mitochondria (purple) and ER-like organelles (green), respectively, are zoomed in and arranged from the left (TEM) to the middle (iDPC-STEM) and from top (mitochondria) to bottom (ER-like organelles). The regions marked by red rectangles in iDPC-STEM images are further enlarged and shown at the right accordingly. In these enlarged images, the structure models of ATP synthase (EMDB entry EMD-11368) can be comfortably fit into the protein density on the mitochondrial cristae membrane (top right) and the membrane protein complexes (marked by the red arrow, bottom right) can be visualized in the luminal space of the ER-like organelle.

For quantitative comparison, a line profile across the mitochondrial membrane has been produced and the contrast calculated within each micrograph as explained in Methods and Figure S3. The results are summarized in Figure 2b. With increase of the electron dose the micrograph contrast of both TEM and iDPC-STEM imaging increases. However, at corresponding doses, the iDPC-STEM contrast stays constantly above (around twice) compared to the conventional TEM contrast, significant even at the extreme low dose value of 1 e/Å^2^.

Beside the enhanced contrast, iDPC-STEM resolves more structural information in comparison with TEM. We further investigated a pair of micrographs in Figure 2a, taken at the relative high exposure dose of 40 e/Å^2^, where TEM should behave optimally. The region of one mitochondrion (purple rectangle) and the region of one mitochondrion-contacted endoplasmic reticulum (ER)-like organelle (green rectangle) in each micrograph were enlarged in Figure 2c. We found that a pair of ATP synthase are resolved on the cristae membrane in the iDPC-STEM micrograph while we could not identify them in the TEM micrograph. In addition, we could observe two membrane protein complexes located at the luminal side of the membrane (red arrow) of the ER-like organelle, while they could not be resolved in the corresponding TEM micrograph.

To prove consistency and reproducibility, we repeated the electron dose experiment for another HepG2 cell specimen. This reconfirmed the results (Figure S4).

### iDPC-STEM contrast vs. TEM contrast with respect to staining

Biological resin sections contain mainly light elements, providing a limited contrast in normal TEM. Therefore, heavy metal (Os, U and Pb) staining has always been used to enhance the contrast. Nevertheless, this staining procedure is complicated, involves human safety concerns, and also, with heavy staining, introduces presumable structural artifacts. Therefore, the future imaging technique is demanded to operate with less staining, while preserving and better resolving the ultrastructure. In our second experiment, we investigate iDPC-STEM in this respect, if it could generate images with good enough contrast for the sections less stained.

We tested two kinds of sections of HepG2 cell, one regularly prepared, using Method 1, and another with the minimal heavy atom staining, using Method 2. The thickness of the sections was again 70 nm. The total electron dose for both TEM and iDPC-STEM was set to 3 e/Å^2^. This time, we selected two different convergence semi-angles 6.43 mrad and 2.58 mrad for iDPC-STEM imaging. The corresponding depths of focus are 121 nm and 751 nm, respectively (Table 1). These are both larger than the thickness of the section and, therefore, the first imaging mode of STEM, shown in Figure 1b, is taking place. The original pixel size of iDPC-STEM images was set to 1.43 nm and then resized to match the pixel size of TEM images by FFT cropping algorithm. For TEM imaging, we set up an appropriate magnification and the defocus value of ∼ 7 μm to image the same region of the section with a similar field of view and the final pixel size on the camera was 1.50 nm.

Comparing the sections prepared with Method 1 (high staining) and Method 2 (minimal staining), we found that the contrast of TEM micrograph significantly decreased with the reduction of heavy atom staining. This is shown qualitatively in Figures 3a, 3b and 3c and quantitatively, by line profile measurement in Figure 3d. We found at the minimal staining condition (Method 2) the SNR of TEM micrograph is quite low. However, at the same time, the reduction of contrast in iDPC-STEM micrograph is much less, both observed qualitatively (Figures 3a, 3b**, and** 3c) and quantitatively (Figure 3d).

**Figure 3.**
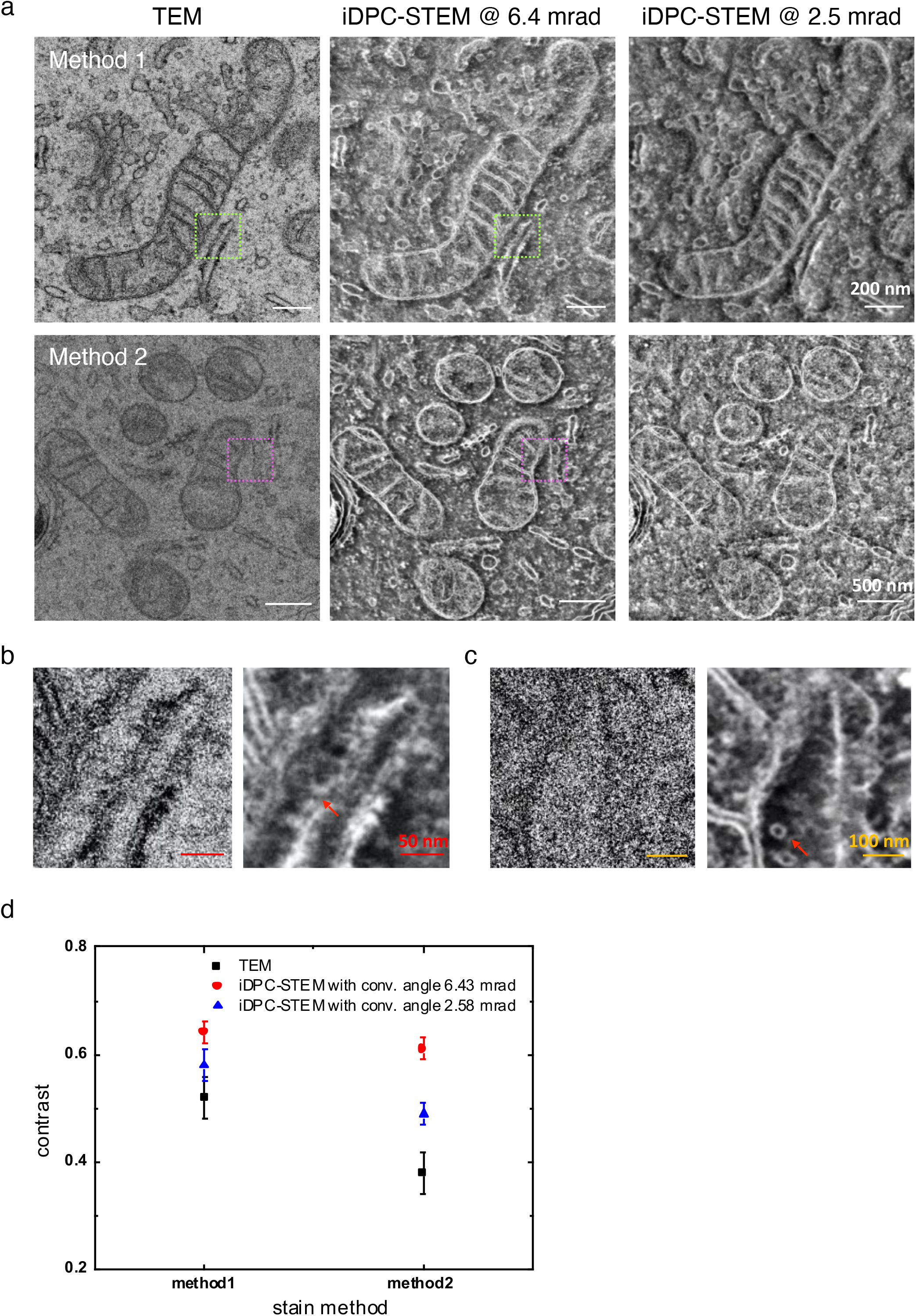
Comparison between TEM and iDPC-STEM imaging for different amount of staining. (**a**) The sections of HepG2 cell prepared using Method 1 and Method 2, respectively. The thickness of the section is 70 nm. TEM images acquired with the defocus of ∼ 7 μm and the total exposure dose of 10 e/Å^2^. The iDPC-STEM images acquired with the total exposure dose of 10 e/Å^2^ and two convergence semi-angles 6.43 mrad and 2.58 mrad, respectively. The pixel size of TEM is 1.50 nm. The original pixel size of iDPC-STEM was 1.43 nm, resized to 1.50 nm by FFT cropping algorithm. (**b**) Regions marked by the green rectangles are zoomed in and compared. Left, TEM. Right, iDPC-STEM. The ribosome particles on the ER membrane are indicated by the red arrow in the iDPC-STEM image. (**c**) Regions marked by the purple rectangles are zoomed in and compared. Left, TEM. Right, iDPC-STEM. The interesting ultrastructures (vesicle or protein complex compartment) between mitochondrion and ER are indicated by the red arrow in the iDPC-STEM image. (**d**) Quantitative comparison of contrast between TEM and iDPC-STEM images for different stained sections. The contrast of each micrograph was measured using the line profile method (see Figure S3). For each data point, the measurement was repeated three times using different profile lines at different regions.

For the section prepared with Method 1 (heavy staining), in consistency with the previous experiment, the contrast of iDPC-STEM at 6.43 mrad is better than in TEM (Figure 3d) and the ribosomes on the surface of rough ER are better resolved in comparison with the corresponding TEM micrograph (indicated with the arrow in Figure 3b). Furthermore, for the section prepared with Method 2 (minimal staining), while TEM contrast deteriorated significantly (Figure 3d), the contrast of iDPC-STEM at 6.43 mrad micrograph was still strong enough to resolve small vesicles (or protein complex compartments) at the cytosolic space between the mitochondria and the contact ER (indicated with the arrow in Figure 3c). This could by no means be observed in the corresponding low contrast (low SNR) TEM micrograph (Figure 3c).

As mentioned, the two different convergence semi-angles (Figure 3a) have been applied for iDPC-STEM in this experiment. As expected from theoretical consideration (Table 1), because sample was thinner than the depth of focus in both cases, the larger convergence semi-angle produced sharper image. The smaller the convergence semi-angle, the larger the STEM probe becomes, resulting in reduced resolution and therefore sharpness (Figure 3d). At reduced resolution, resolving the double membrane feature of mitochondria (Figure 3a) naturally becomes harder, in some cases, worse than TEM. Therefore, selection of a proper convergence semi-angle is important for iDPC-STEM technique. Whenever sample allows (e.g. thinner) larger convergence angle should be used to enhance resolution and sharpness.

To prove consistency and reproducibility, we repeated the staining experiment for another HepG2 cell specimen. This reconfirmed the results (Figure S5).

### iDPC-STEM vs. TEM contrast and structural details for thick sections

One of the important advantages of iDPC-STEM imaging is the ability to image thicker samples by applying smaller convergence angle of the beam (Bosch and Lazić, 2019). The smaller convergence angle of the beam provides a larger depth of focus, that can easily become larger than the sample thickness. This enables the full projection of the sample electrostatic potential and enhancing the contrast for all the features extending along the sample in-depth direction.

In the next experiment, the section samples of mice testis tissues of different thicknesses, i.e., 200 nm, 400 nm and 600 nm, prepared using Method 1, have been investigated. The total exposure dose for both TEM and iDPC-STEM was set to 10 e/Å^2^. Two different convergence semi-angles 6.43 mrad and 2.58 mrad for iDPC-STEM imaging have been used, corresponding depths of focus as 121 nm and 751 nm, respectively (Table 1). The former is shorter, while the latter is longer than the thickness of the sections used in the experiment. The pixel size of iDPC-STEM images was set to 1.1 nm. For TEM imaging, we set up an appropriate magnification and the defocus value of ∼ 7 μm to image the same region of the section with a similar field of view and the final pixel size on the camera was 1.04 nm.

For the 200 nm thick section, conventional TEM still produces sharp image with a reasonable contrast (Figure 4a, left). The current thickness is still within the mean free path of biological section for 200 keV electron, which was measured to be 300 nm (Yan et al., 2015). However, the effect of the focus gradient at the current thickness cannot be avoided (Sun, 2018), affecting the resolution and contrast in TEM. For iDPC-STEM both at 6.43 mrad and 2.58 mrad (Figure 4a, middle and right, respectively), good quality micrographs have been produced, as good as the ones for 70 nm sections (Figure 2a). A better contrast in comparison with the TEM image has been obtained, qualitatively as well as qualitatively (see Figure 4d). Due to comparable depth of focus at 6.43 mrad (121 nm) with the sample thickness (200 nm), the image contrast and resolution of iDPC-STEM at 6.43 mrad is better than that of iDPC-STEM at 2.58 mrad.

**Figure 4.**
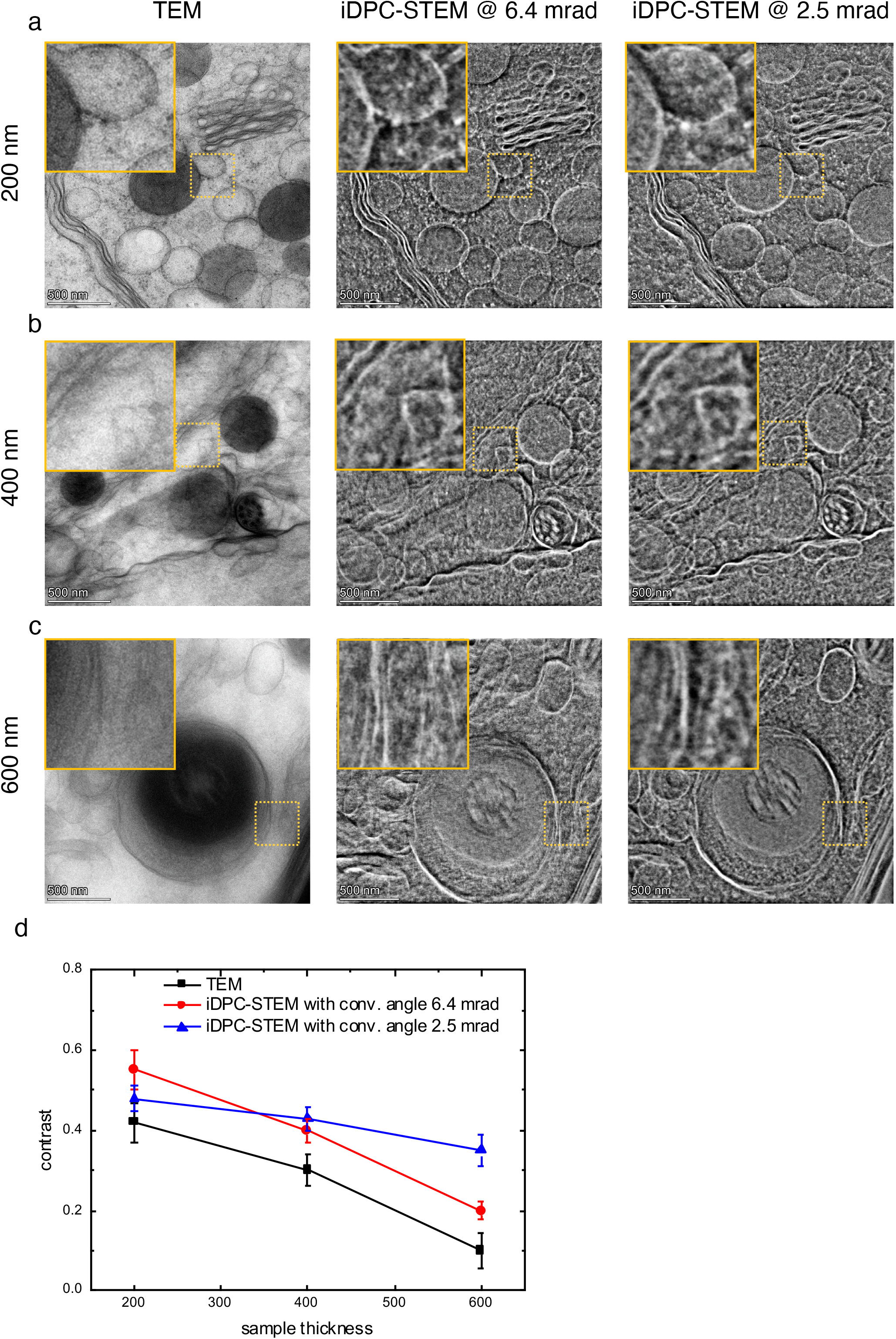
Comparison between TEM and iDPC-STEM imaging for different section thicknesses. The sections of mice testis tissues were prepared using Method 1 with different thickness of 200 nm (a), 400 nm (b) and 600 nm (c), respectively. TEM images (left) were acquired with the defocus of ∼ 7 μm and the total exposure dose of 10 e/Å^2^. The iDPC-STEM images were acquired with the total exposure dose of 10 e/Å^2^ and two convergence semi-angles 6.43 mrad (middle) and 2.58 mrad (right), respectively. The pixel size of iDPC-STEM imaging is 1.1 nm. The original pixel size of TEM images was 1.05 nm and then resized to 1.1 nm by FFT cropping algorithm. (**d**) Quantitative comparison of contrast between TEM and iDPC-STEM images for different thickness of sections. The contrast of each micrograph was measured using the line profile method (see Figure S3). For each data point, the measurement was repeated three times using different profile lines at different regions.

For the section of 400 nm thickness, the quality of TEM micrographs deteriorate, both due to inelastic scattering of electrons and defocus CTF canceling with most of high frequency contrast lost. As the result, many fine membrane structures can no longer be resolved (Figure 4b, left). However, for iDPC-STEM both at 6.43 mrad and 2.58 mrad good contrast micrographs are obtained, resolving many fine structural details of the membrane (Figure 4b, middle and right). Quantitatively this is also confirmed (Figure 4d). As expected, the image contrast of iDPC-STEM at 6.43 mrad became worse than what was obtained for the 200nm thickness. This is because the thickness of 400 nm is now noticeably larger than the depth of focus of 121 nm and the image contrast is influenced by contributions from the regions below and above the central part of the probe (Figure 1b).

For even thicker section, of 600 nm thickness, the quality of TEM micrograph deteriorates further, for the same reasons as explained above, becoming practically useless (Figure 4c, left). Only the low frequency contrast features, like membrane shapes, are resolved. iDPC-STEM at the other hand, is barely affected and deteriorated only minorly. Besides the shape, detailed membrane ultrastructure is resolved from the micrographs, both at 6.43 mrad and 2.58 mrad. The image contrast obtained at 2.58 mrad by iDPC-STEM is now stronger than the one obtained by iDPC-STEM at 6.43 mrad (see quantitative comparison in Figure 4d). This is because the thickness of the section (600 nm) is now significantly larger than the depth of focus of iDPC-STEM at 6.43 mrad (121 nm), while still smaller than the depth of focus at 2.58 mrad (751 nm).

To prove consistency and reproducibility, we repeated the thickness dependence experiment for another mouse testis tissue specimen. This reconfirmed the obtained results (Figure S6).

### Depth-sectioning of iDPC-STEM imaging for thick samples

When the depth of focus of the electron beam is smaller than the sample thickness (Figure 1b, bottom), besides electron tomography, to resolve the in-depth structural information of a thick sample, a depth-sectioning method can be applied (Bosch and Lazić, 2019; de Jonge et al., 2010). Here, we demonstrate this capability using iDPC-STEM for biological sample. Although not optimal (ideally should be large > 30 mrad), a convergence semi-angle of 6.43 mrad used here has been good enough to show some structural variation at different depths.

We used Method 1 to prepare a resin embedded section of another mouse testis tissue with the thickness of 600 nm. The total exposure dose for iDPC-STEM was set to 3 e/Å^2^. The pixel size of iDPC-STEM images was set to 5 nm. We acquired a series of iDPC-STEM images using different probe focal positions from over-focus (above the sample) to under-focus (below the sample). The defocus step was chosen as 10 nm. From this stack of the images, slices separated by 50 nm, representing the middle part of the sample have been selected and shown in Figure 5. Zero defocus is set to be in the middle of the sample. By investigating the slices, we found that the membrane structure regions change at different depths. We indicated these regions in Figure 5 showing that the depth-sectioning did have an effect.

**Figure 5.**
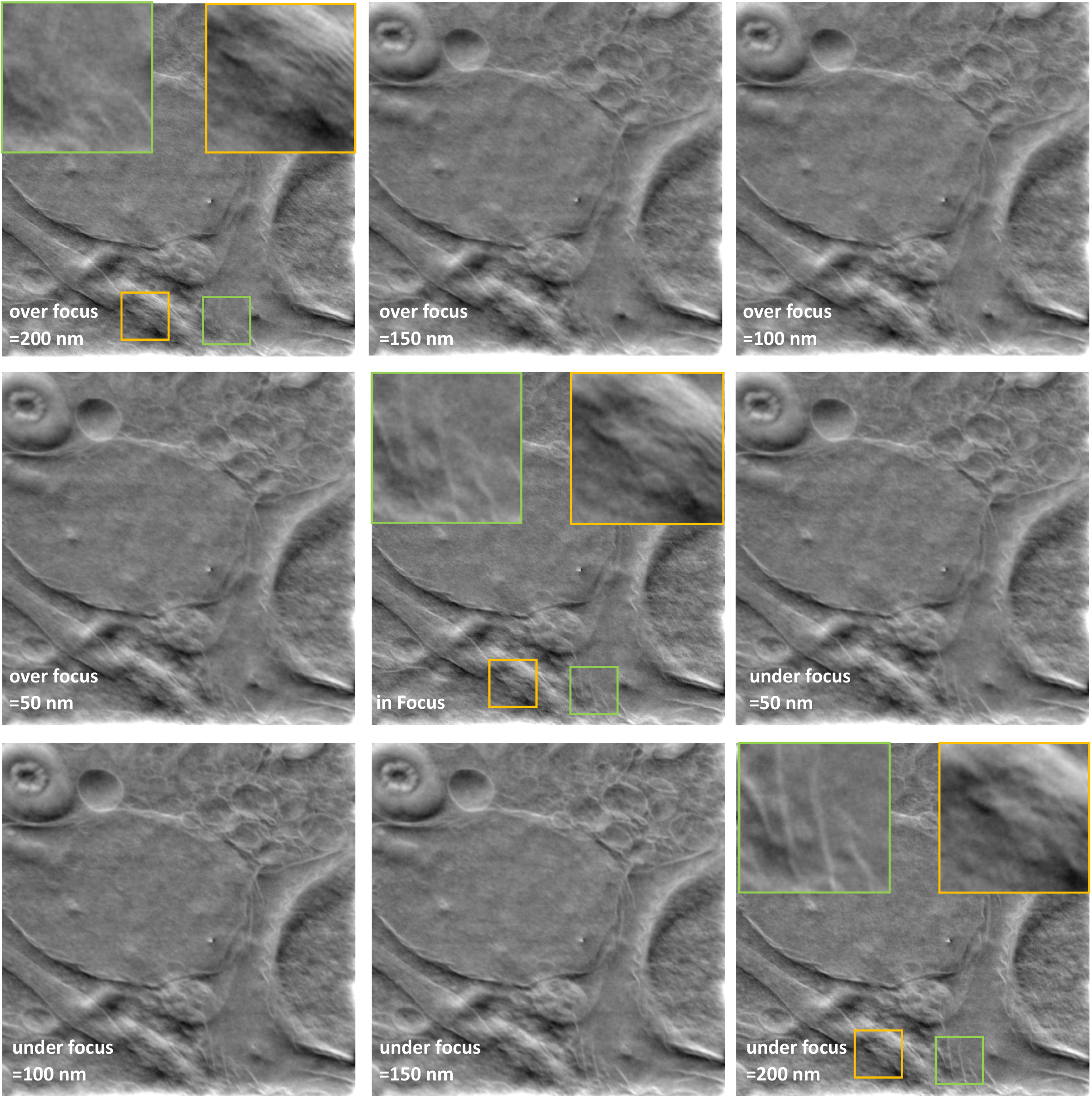
Depth-sectioning using iDPC-STEM imaging for a thick section. The resin embedded mouse testis tissue was prepared using Method 1 and sectioned to thickness of 600 nm. Series of iDPC-STEM images were acquired with convergence semi-angle of 6.43 mrad and the exposure dose of 3 e/Å^2^. The pixel size was set to 5 nm. The defocus step between selected images is 50 nm. The central slices covering almost the full thickness of the sample area are shown sequentially from top to bottom with the focus positions centered in the middle of the sample.

Note that the depth of focus with this convergence semi-angle is still quite large (121 nm), meaning the in-depth resolution is very low. Therefore, revealing the full 3D structural information is certainly not to be expected. Appropriate depth-sectioning can be provided with an aberration corrected probe, formed with the probe aberration corrected system (Borisevich et al., 2006). This allows larger convergent semi-angles, producing the probes smaller in all 3 directions (Figure S1, top row).

## DISCUSSIONS

Attempt to apply STEM technology into biological research has been going on for almost half a century, from molecular mass measurement (Colliex and Mory, 1994) to tomography of biological sections (Aoyama et al., 2008; Yakushevska et al., 2007). However, compared to TEM, the previously available STEM techniques had not shown a significant advantage. In the present work, we investigated, recently emerged STEM technique, iDPC-STEM (Lazić et al., 2016; Yucelen et al., 2018), which could solve the low dose-efficiency issue of (HA)ADF-STEM and the non-linear contrast problem of (A)BF-STEM.

Considering the significant radiation damage effect of biological specimen, we first studied whether iDPC-STEM could yield good contrast of biological sections with limited dose. We found that, from normal low dose condition of 40 e/Å^2^ to extreme low dose condition of 1 e/Å^2^, iDPC-STEM could always generate reasonable contrast better than conventional TEM. When the dose is below 3 e/Å^2^, discerning membrane ultrastructure becomes difficult in TEM, while it is still clear in iDPC-STEM. We consider two main reasons to explain this observation.

First, the vector data integration step in iDPC-STEM naturally reduces the noise, as noise is a non-conservative vector field, while fully preserving the conservative electric field vector which is our primary signal (Lazić et al., 2016). This increases signal-to-noise ratio (SNR) of the final micrograph. The effect of adding the noise to corresponding DPC-STEM components and the noise transfer to the final iDPC-STEM image is demonstrated in Figure S7. Even when most of the signal spectrum is drown into noise in the component images (SNR ≤ 1), most of the spectrum stays above noise (SNR ≥ 1) in the final image. Such enhancement could not appear within TEM nor any other conventional STEM technique.

Second, the contrast transfer function oscillations and low frequency transfer. For heavy metal-stained resin-embedded section, as reported before, the contrast in TEM micrograph is majorly contributed from the electron exit wave amplitude (Reimer and Kohl, 2008). The electron wave phase information of resin-embedded section contributing from the cross-section difference between biological material and resin is small (Carlemalm and Kellenberger, 1982) and significantly faded by CTF oscillation effect in TEM. However, for iDPC-STEM, with positive definite CTF and full frequency transfer including low frequencies (Lazić et al., 2016) the result is similar to the effect of using phase plate in TEM (Danev and Nagayama, 2001; Danev et al., 2009; Edgcombe, 2016). This has been proven earlier (Lazić and Bosch, 2017) and in our experimental result shown in Figure 2. For the imaging result at 40 e/Å^2^, good contrast is observed for stained membrane structure in both TEM and iDPC-STEM, while the less stained membrane protein complexes could be only visualized using iDPC-STEM.

Besides dose efficiency, we investigated iDPC-STEM imaging of biological sections with the minimal staining. Since phase information is fully transferred in iDPC-STEM, a good contrast for less stained biological sections is expected. Our experimental data showed that while at the minimal stain condition the contrast of TEM micrograph significantly deteriorates iDPC-STEM micrographs do show good contrast and ultrastructural details at molecular level (Figure 3). This observation encouraged us to try iDPC-STEM technique on the unstained resin-embedded biological section, which was previously studied by HAADF-STEM (Carlemalm and Kellenberger, 1982). More interestingly, applying iDPC-STEM technique on the cryo-vitrified lamella of biological specimen would be a promising direction in the future.

Finally, we studied the influence of sample thickness on iDPC-STEM and TEM. The conventional TEM quality deteriorates rapidly with poor contrast with increase of sample thickness. There are two reasons for this phenomenon. The focus gradient resulting cancellation of CTF as the major one (Sun, 2018) and the significantly increased portion of inelastically scattered and multiple scattered electrons that contribute to the image background. The second becomes significant when the sample thickness is beyond the mean free path of accelerated electron (Egerton, 2011). Our experimental results confirm these effects. For the sample thickness of 200 nm, within the mean free path of 200 keV electron (Yan et al., 2015), the TEM micrograph still yields good contrast with a bit reduced structural details, due to focus gradient. When the sample thickness increases to 400 nm and 600 nm, this reduction becomes significant, for most of the features, terminal.

For iDPC-STEM, when a proper semi-convergence angle is selected, the depth of focus can be large enough to cover the whole range of the sample thickness, thus avoiding the focus gradient problem. This is due to amorphous like nature of biological samples that do not significantly alter and disturb the probe shape (Bosch and Lazić, 2019). As a result, iDPC-STEM is suitable to image thick samples, which we confirm in our experimental investigation. With the convergence semi-angle of 2.5 mrad that yields a depth of focus of 751 nm, the iDPC-STEM image contrast is constantly high for all sample thickness cases (200nm, 400nm and 600nm), preserving detailed membrane ultrastructure in the micrographs. This is because the 751 nm depth of focus is comfortably larger than all these the sample thicknesses (Figure 1b, top row). With the convergence semi-angle of 6.43 mrad that yields a depth of focus of 121 nm, the contrast of iDPC-STEM is well preserved for the sample thickness of 200 nm (comparable to 121 nm) while becoming slightly worse for the thicknesses of 400 nm and 600 nm (Figure 4d). This is due to depth of focus becoming smaller and smaller with respect to sample thickness (Figure 1b, bottom row). In this case, only part of the central probe waist-length provides sharp signal of interest, while parts of the sample above and below the beam waist interact with defocused parts of the probe and contribute to the random-like background signal. This is what causes a drop in contrast, although offers an opportunity of depth-sectioning. Note that, even though at certain point high convergence semi-angle iDPC-STEM contrast becomes smaller than the contrast obtained with small convergence semi-angle, it is still significantly better than the contrast obtained with TEM.

In iDPC-STEM, a smaller convergence semi-angle needed to yield large enough depth of focus, decreases the achievable in plane resolution (Table 1). Therefore, to preserve high resolution for thick sample, a technique of depth-sectioning would have to be performed (Bosch and Lazić, 2019). In the present work, we performed only a proof-of-concept experiment using the semi-convergence angle of 6.43 mrad and depth of focus of 121 nm (Figure 5). This depth of focus is still large and cannot yield a good resolution in the vertical direction. To achieve better resolution in vertical direction, there are two possible ways. One is to further increase the semi-convergence angle and reduce the depth of focus, which can be only performed in a Cs-corrected microscope. Another way is to combine depth sectioning and tomography to improve the resolution in both lateral and vertical directions, which has been already reported for BF-STEM (Trepout et al., 2015). This is out of the scope of this work.

Overall, we would like to conclude that iDPC-STEM technique shows a high performance for studying resin-embedded biological section by providing fruitful contrast in low dose imaging condition, suitable for minimal stains and owning unique advantages for hick sample (Table 2), implying a promising future for its wide application in the biological research.

**Table 2.** A summary of comparison between TEM and iDPC-STEM imaging in different experiments.

|  | TEM<sup>1</sup> | iDPC-STEM |
| --- | --- | --- |
| Contrast of samples in varies dose:<br>Total dose < 10 e/ Å<sup>2</sup><br>Total dose > 10 e/ Å<sup>2</sup> | Medium<br>High | High<br>Very high |
| Contrast of samples with varies amount of stain:<br>Method 1 (normal OsO4 + post stain)<br>Method 2 (normal OsO4) | High<br>Medium at<br>membrane area<br>but weak at<br>protein area | High<br>High at both<br>membrane and<br>protein area |
| Contrast of samples with varies thickness:<br>200nm<br>400nm<br>600nm | High<br>Medium<br>Weak | High<br>High<br>High<sup>2</sup> |
| Depth sectioning | Not available | Possible<sup>3</sup> |

<sup>1</sup> all TEM images were taken on FEI Talos F200C with an FEI Ceta camera.
<sup>2</sup> convergence semi-angles of 2.58 mrad
<sup>3</sup> convergence semi-angles of 6.43 mrad

## AUTHOR CONTRIBTUIONS

F. S. and I. L. initiated and supervised the project. X. L. and X. H. performed all microscopy experiments. L. W. prepared resin embedded sections. X. L., X. H., I. L. and Y. D. analyzed experimental data. D. W, M. W. and L. Y. contributed to discussion of experimental design and data analysis. X. L., I. L. and F.S. wrote the paper.

## ACKNOWLEDGMENTS

We would also like to thank Yanxia Jia (CBI, IBP) for her assistance of project management. This work was equally supported by grants from National Natural Science Foundation of China (31830020), the Ministry of Science and Technology of China (2017YFA0504700) and Chinese Academy of Sciences (XDB37040102). The authors would also like to thank the grant supports from Beijing Municipal Science and Technology Commission (Z181100004218002). All the sample preparation and electron microscopy work were performed at Center for Biological Imaging (CBI, http://cbi.ibp.ac.cn), Institute of Biophysics, Chinese Academy of Sciences.

## COMPLIANCE WITH ETHICAL STANDARDS

All authors declare that they have no conflict of interest. All institutional and national guidelines for the care and use of laboratory animals were followed.

## SUPPLEMENTARY FIGURE LEGENDS

**Figure S1.**
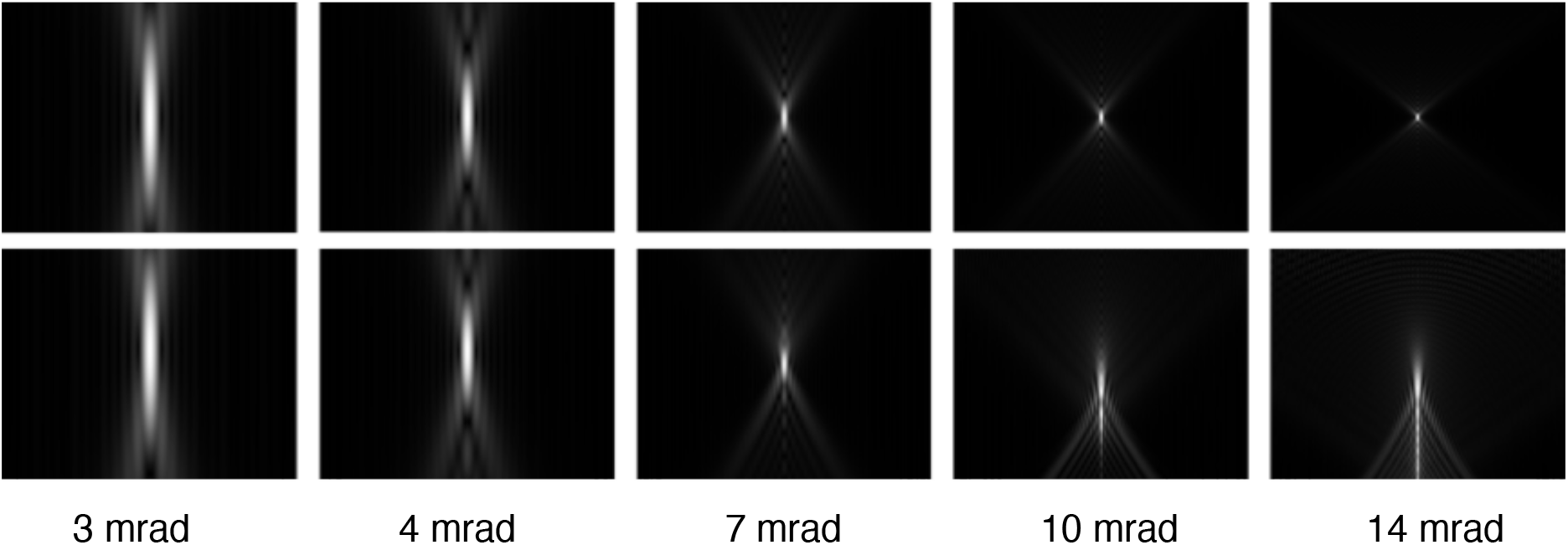
Theoretical vertical cross section of the probe along the direction of 200keV electron beam propagation (from top to bottom) with different convergence semi-angles. Top row, probe corrected system (Cs=0). Bottom row, probe non-corrected system (Cs = 2.7 mm). For the non-corrected system, the critical semi-angle at 7.4 mrad where the probe reaches the smallest size. The cross section was straightforwardly generated using Fresnel propagator (or equivalently defocusing) starting from the focus position of the condenser system (Kirkland, 2010).

**Figure S2.**
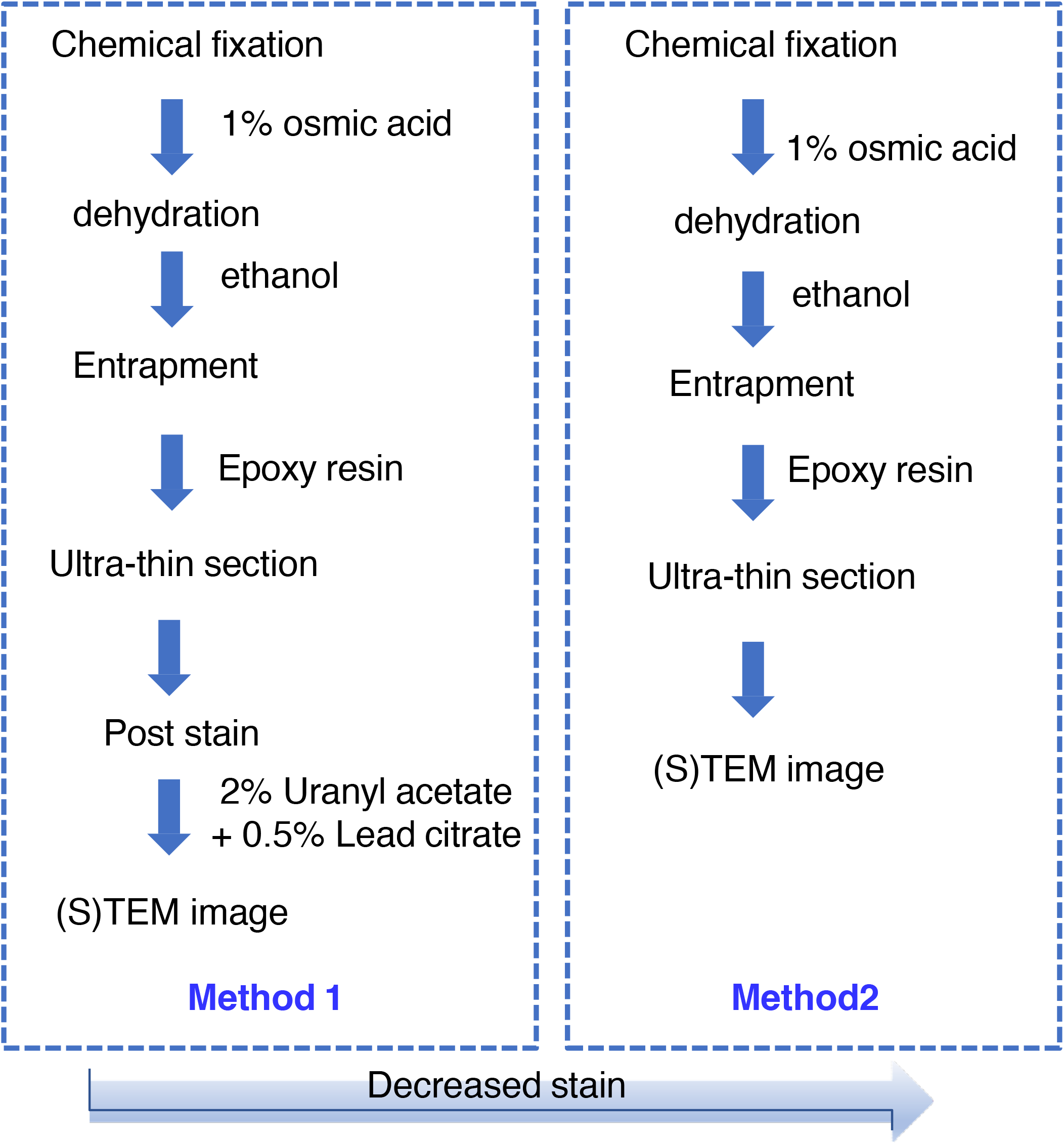
Sample preparation procedures of Method 1 and Method 2 (see also Materials and Methods).

**Figure S3.**
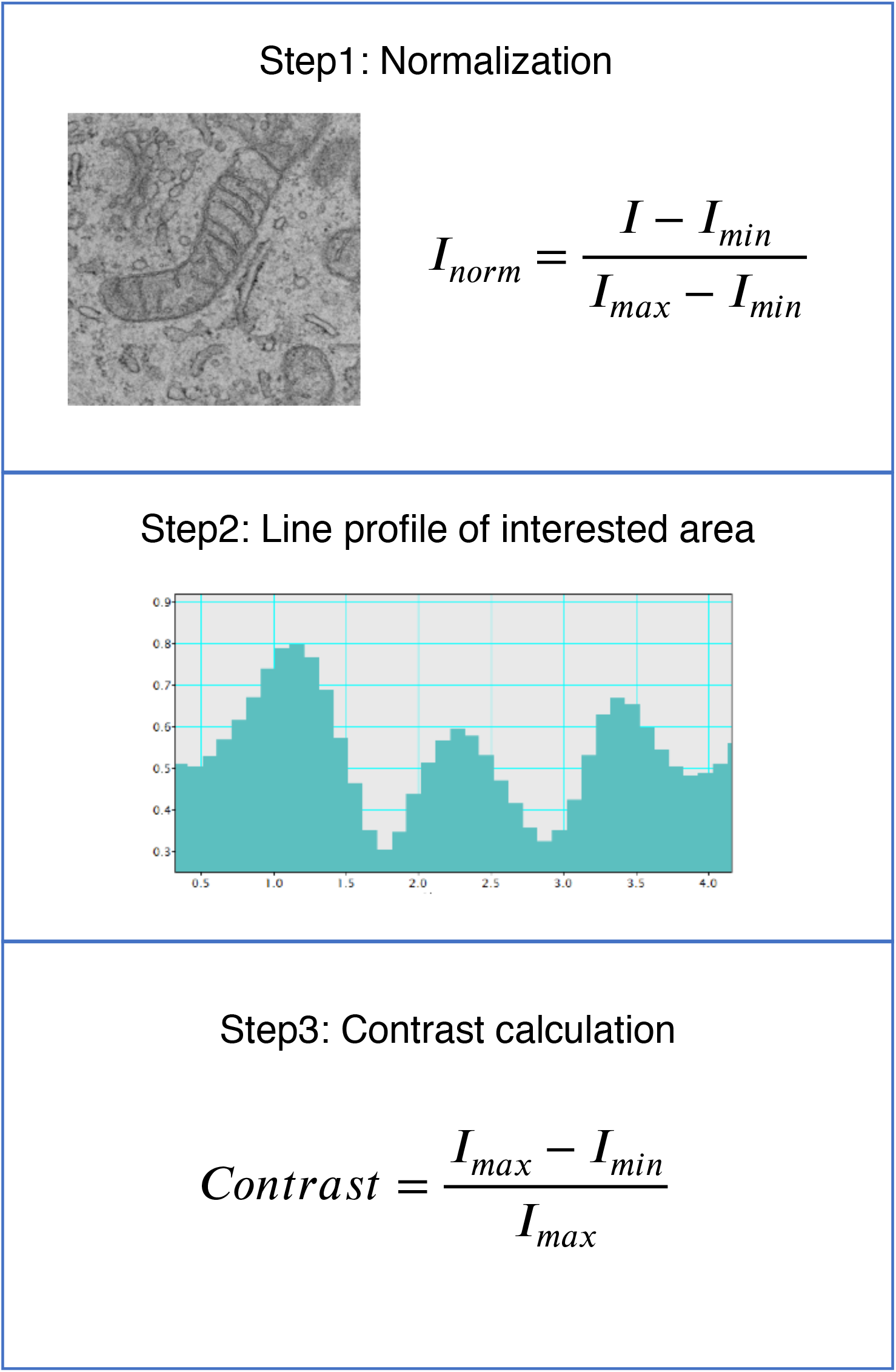
The contrast measurement by using line profile method. Step 1, the images normalization according to the pixel variance. Step 2, the line profile generation at the region of interest. Step 3, the contrast computation as difference between maximum and minimum intensity along the profile, normalized with the maximum.

**Figure S4.**
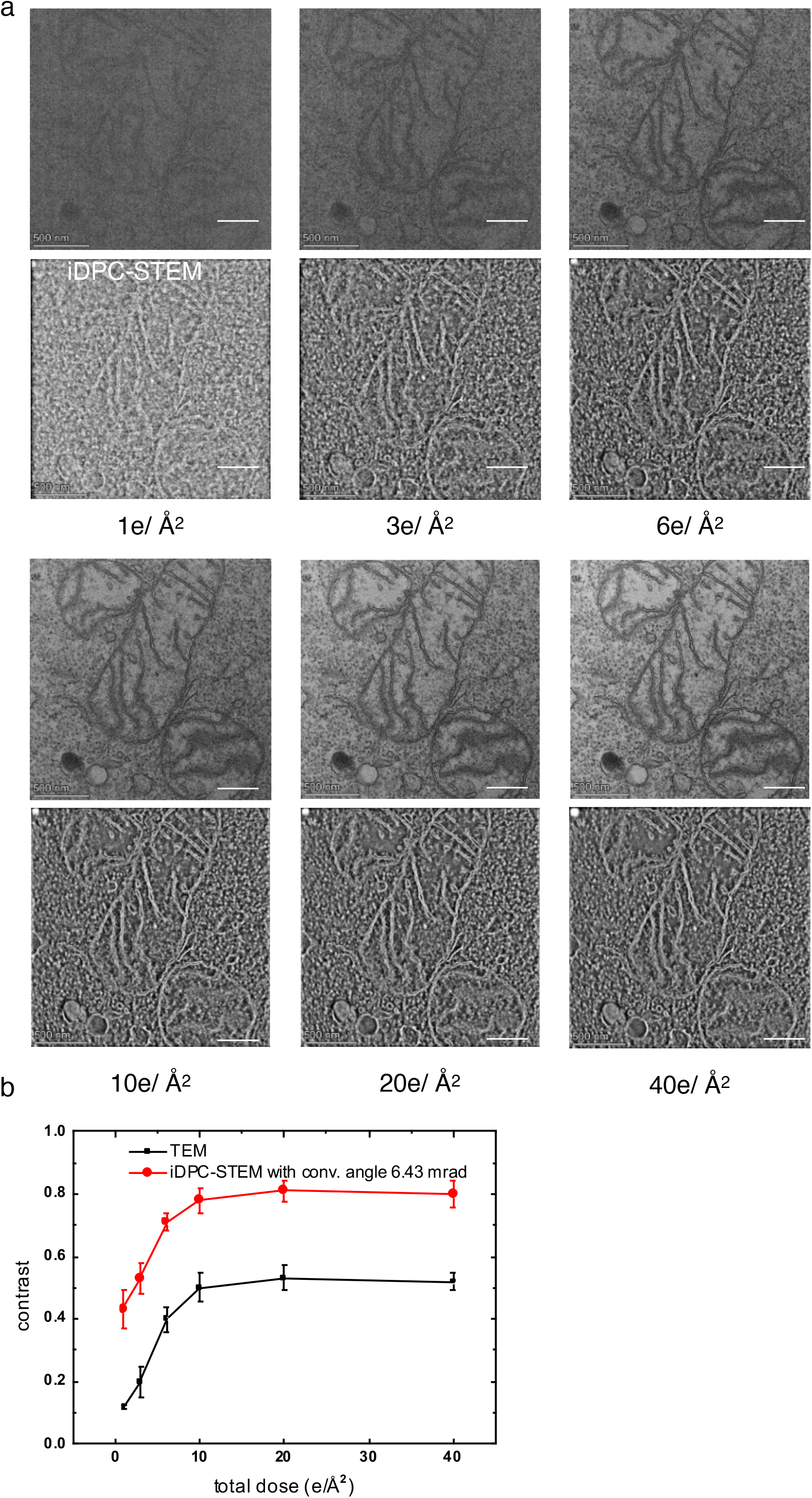
Comparison between TEM and iDPC-STEM imaging at different electron doses for another section of HepG2 cell. (**a**) TEM and iDPC-STEM micrographs acquired with the total electron doses of 1 e/Å^2^, 3 e/Å^2^, 6 e/Å^2^, 10 e/Å^2^, 20 e/Å^2^ and 40 e/Å^2^, respectively. The pixel size TEM imaging is 1.04 nm. The pixel size of iDPC-STEM images was 1.1 nm. The convergence semi-angle of iDPC-STEM is 6.43 mrad. (**b**) Quantitative comparison of contrast between TEM and iDPC-STEM images at different exposure doses. The contrast of each micrograph measured using the line profile method (see Figure S3). For each data point, the measurement was repeated three times using different profile lines at different regions.

**Figure S5.**
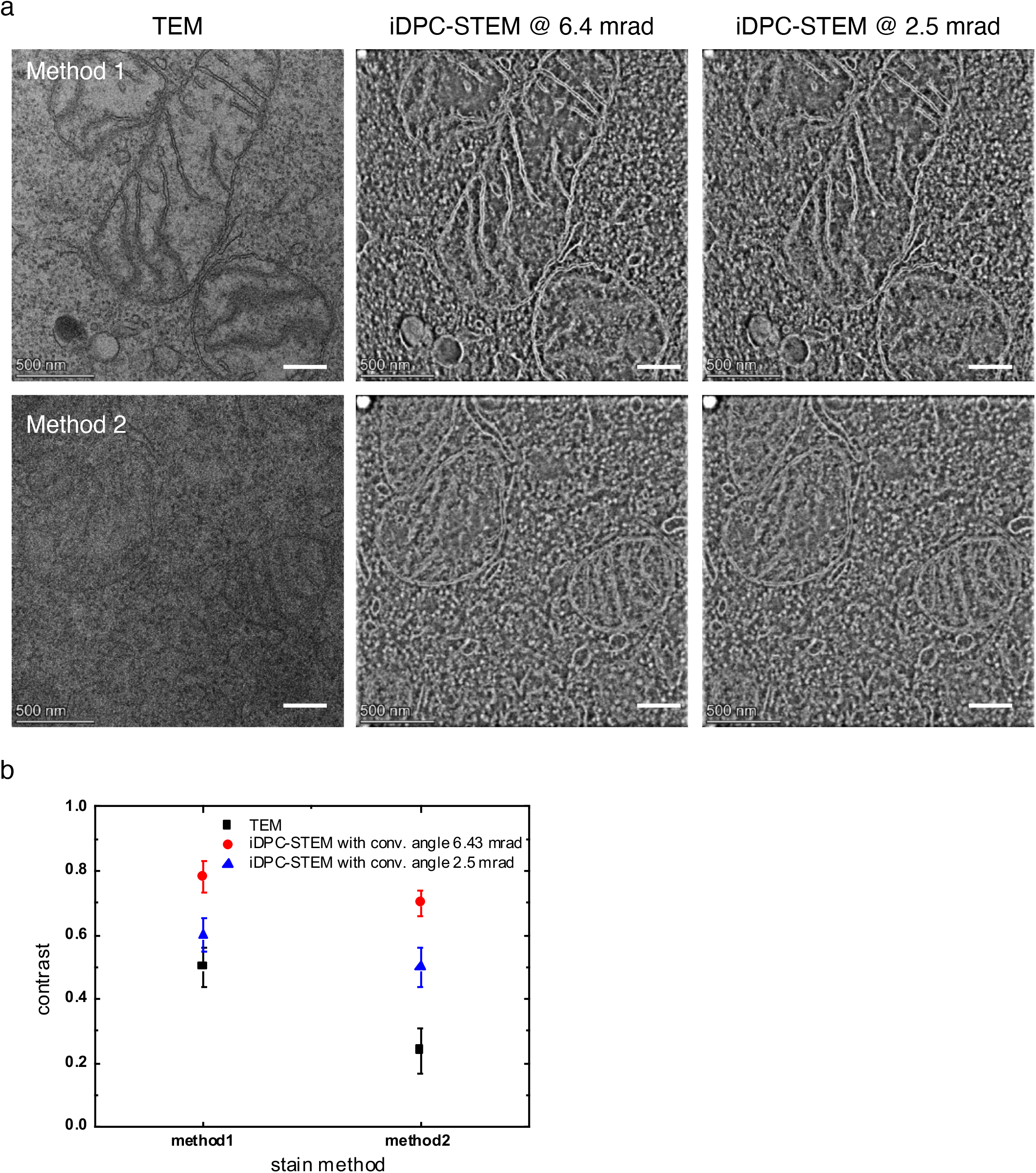
Comparison between TEM and iDPC-STEM imaging for different amount of staining using another section of HepG2 cell. (**a**) The sections prepared using Method 1 and Method 2, respectively. The thickness of the sections is 70 nm. TEM images were acquired with the defocus of ∼7 μm and the total exposure dose of 10 e/Å^2^. The iDPC-STEM images were acquired with the total exposure dose of 10 e/Å^2^ and two convergence semi-angles 6.43 mrad and 2.58 mrad, respectively. The pixel size of TEM imaging is 1.04 nm. The pixel size of iDPC-STEM images was 1.1 nm. (**b**) Quantitative comparison of contrast between TEM and iDPC-STEM images for different stained sections. The contrast of each micrograph measured using the line profile method (see Figure S3). For each data point, the measurement was repeated three times using different profile lines at different regions.

**Figure S6.**
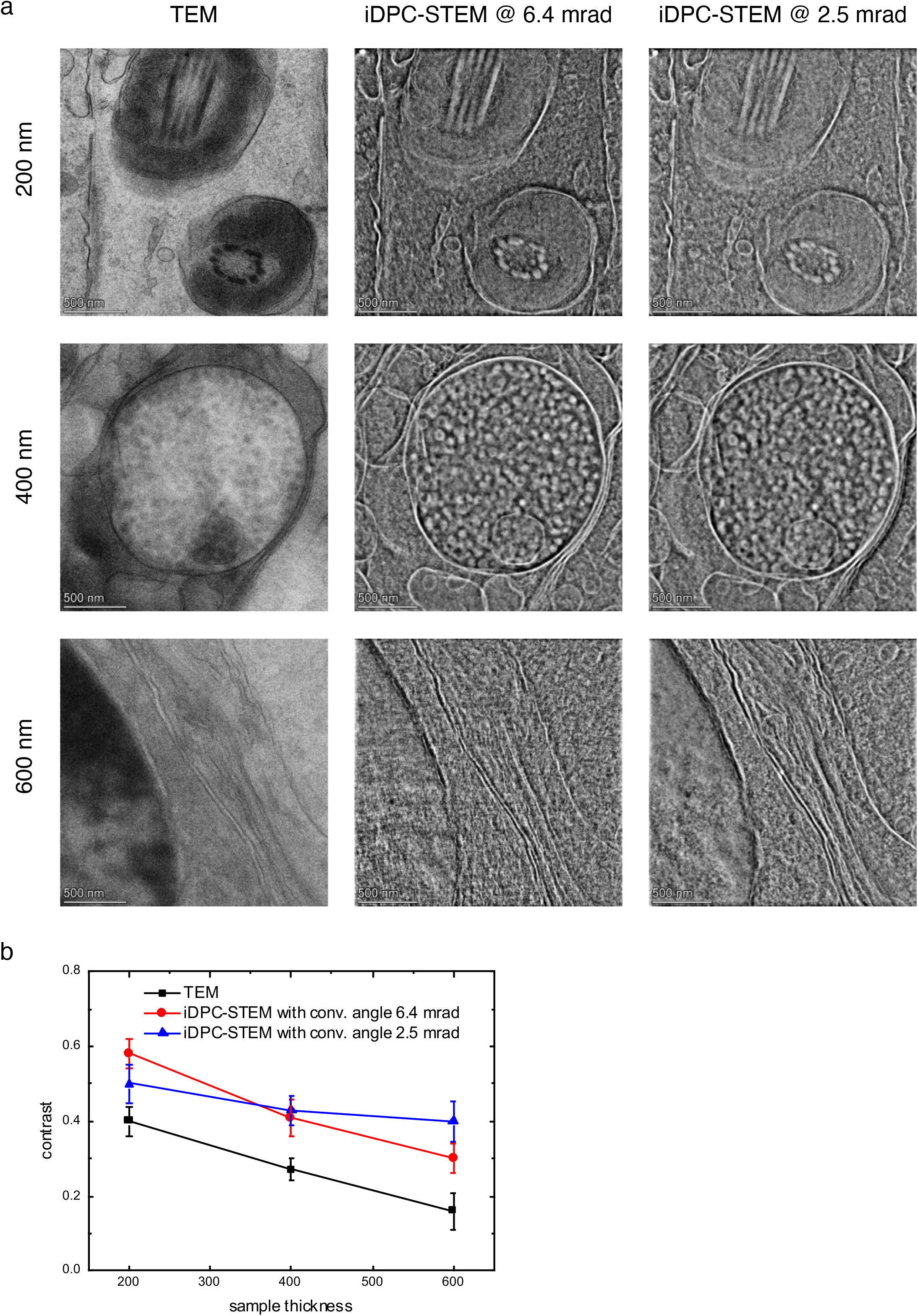
Comparison between TEM and iDPC-STEM imaging upon different section thicknesses using another testis tissue specimen. **(a)** The sections were prepared using Method 1 with different thickness of 200 nm, 400 nm and 600 nm, respectively. TEM images (left) were acquired with the defocus of ∼ 7 μm and the total exposure dose of 10 e/Å^2^. The iDPC-STEM images were acquired with the total exposure dose of 10 e/Å^2^ and two convergence semi-angles 6.43 mrad (middle) and 2.58 mrad (right), respectively. The pixel size of iDPC-STEM imaging is 1.1 nm. The pixel size of TEM images was 1.04 nm. (**d**) Quantitative comparison of contrast between TEM and iDPC-STEM images for different section thicknesses. The contrast of each micrograph measured using the line profile method (see Figure S3). For each data point, the measurement was repeated three times using different profile lines at different regions.

**Figure S7.**
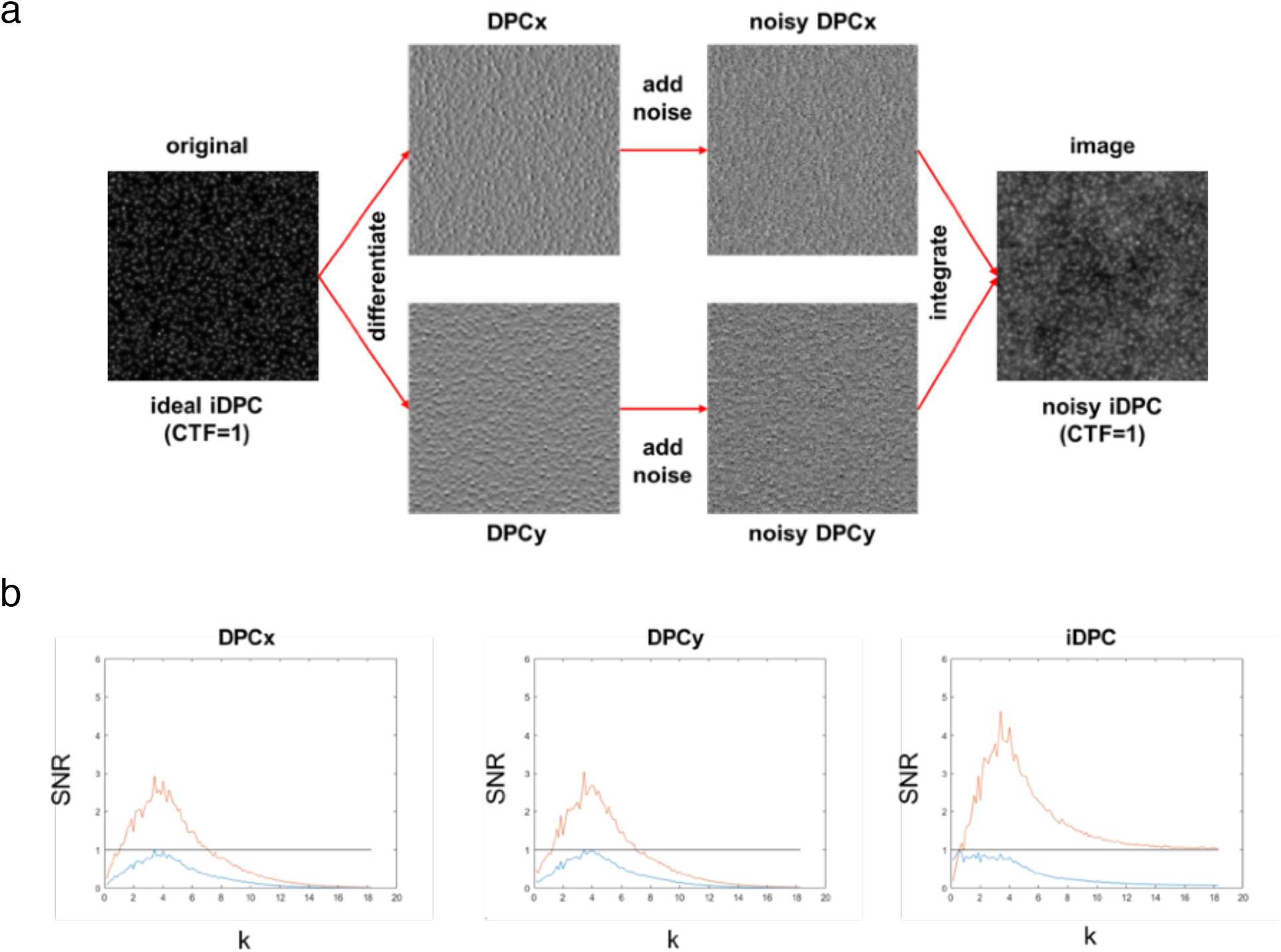
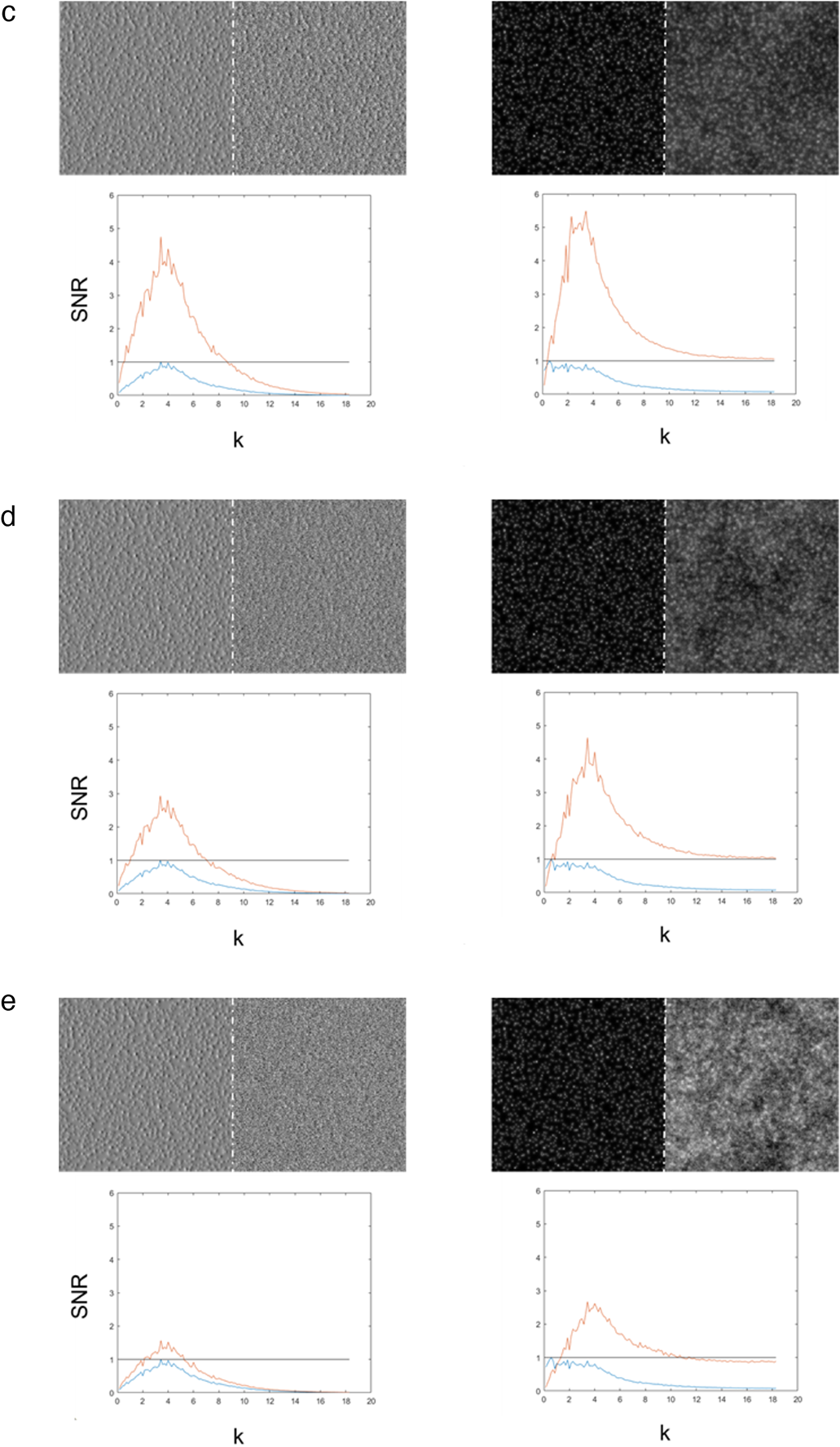
Noise reduction effect by integration step in iDPC-STEM. (**a**) Noise propagation into iDPC-STEM image. Starting from hypothetical, ideal (CTF=1), noiseless iDPC-STEM image, the two gradient components (differentials, DPCx and DPCy) are formed (ideal, noiseless DPC-STEM images). In practice, the two DPC-STEM components acquired directly by measurement are always noisy. Therefore, Gaussian noise is added and inverted gradient operator (integration) is applied (Lazić et al., 2016) to form the noisy iDPC-STEM image. (**b**) Comparison of spectrum signal-to-noise ratio (SSNR, red curves) between obtained noisy DPC-STEM (x and y direction) image and final iDPC-STEM image. Because ideal noiseless images are available, the SSNR is computed directly by dividing the spectrum of the noiseless image (signal) with the spectrum of the difference (noise) between the noisy image and the noiseless image. The spectra of the noiseless images are shown for reference (blue curves), each normalized to corresponding maximum value. As we can see, even though amount of noise added to DPCx/DPCy is such to produce SSNR of 2.7 at the maximum spectral component, the resulting SSNR for iDPC-STEM image is very close to 4 at the same frequency. Clearly, iDPC-STEM reduces noise and increases SSNR. (**c**) (**d**) (**e**) More comparison of SSNR between DPC-STEM and iDPC-STEM images with increasing noise levels. The Gaussian noise added to DPC-STEM to result in SSNR at the spectral maximum reaching (c) 4, (d) 2.7 and (e) 1.4, respectively. Corresponding iDPC-STEM maximum SSNR reached (c) 5, (d) 4 and (e) 2.7, respectively. Left (c-e): the DPCx-STEM component without/with Gaussian noise (top) and the corresponding SSNR curve (bottom). Right (c-e): the corresponding noise-free and obtained noisy iDPC-STEM image (top) and the corresponding SSNR (bottom). **The two main crucial observations: (1)** The SSNR ratio between iDPC-STEM and DPC-STEM increases with more noise added (at maximum (c) 5/4 (d) 4/2.7 and (e) 2.7/1.4). **(2)** Most of the iDPC-STEM SSNR spectrum stays above 1 in all the cases.

